# Structure of a novel α-synuclein filament fold from multiple system atrophy

**DOI:** 10.1101/2024.07.01.601365

**Authors:** Nicholas L. Yan, Francisco Candido, Eric Tse, Arthur A. Melo, Stanley B. Prusiner, Daniel A. Mordes, Daniel R. Southworth, Nick A. Paras, Gregory E. Merz

**Affiliations:** Institute for Neurodegenerative Diseases, University of California San Francisco, San Francisco, CA, USA; Department of Neurology, University of California San Francisco, San Francisco, CA, USA; Department of Biochemistry and Biophysics, University of California San Francisco, San Francisco, CA, USA; Department of Pathology, University of California San Francisco, San Francisco, CA, USA; Department of Pathology, Massachusetts General Hospital, Boston, MA, USA

**Keywords:** Cryo–electron microscopy, α-synuclein, multiple system atrophy, protein aggregation, neurodegeneration, prion

## Abstract

Multiple system atrophy (MSA) is a synucleinopathy, a group of related diseases characterized by the accumulation of α-synuclein aggregates in the brain. In MSA, these aggregates form glial cytoplasmic inclusions, which contain abundant cross-β amyloid filaments. Structures of α-synuclein filaments isolated from MSA patient tissue were determined by cryo–electron microscopy (cryo-EM), revealing three discrete folds that are distinct from α-synuclein filaments associated with other synucleinopathies. Here, we use cryo-EM classification methods to characterize filaments from one individual with MSA and identify a novel, low-populated MSA filament fold (designated Type I_2_) in addition to a predominant class comprising MSA Type II_2_. The 3.3-Å resolution structure of the Type I_2_ filament reveals a fold consisting of two asymmetric protofilaments. One is identical to a previously solved Type I protofilament, while the second adopts a novel fold that is chimeric between two previously reported Type I and II protofilaments. These results further define disease-specific folds of α-synuclein filaments that develop in MSA and have implications for the design of therapeutic and diagnostic molecules that target disease.

## Introduction

Alpha-synuclein is an intrinsically disordered, 140–amino acid protein [7] that localizes primarily to the axon terminals of presynaptic neurons, where it participates in membrane binding, vesicle trafficking, and neurotransmitter release [3, 15]. The misfolding and aggregation of α-synuclein in the brain are hallmarks of a group of neurodegenerative diseases known as synucleinopathies [2, 19], which includes Parkinson’s disease and dementia with Lewy bodies. One such disease, multiple system atrophy (MSA), is clinically characterized by cerebellar dysfunction or parkinsonism and neuropathologically characterized by glial cytoplasmic inclusions (GCIs) in oligodendrocytes [12, 18]. GCIs contain abundant fibrils that appear to be the result of soluble α-synuclein misfolding into self-templating prion conformations that ultimately misassemble as insoluble amyloid filaments [21, 28–31]. Structural studies on filaments isolated from five MSA patients revealed three distinct filament folds (denoted Type I, II_1_, and II_2_). These findings contrast with Lewy body pathologies, which appear to contain a single α-synuclein filament type, and juvenile-onset synucleinopathy, which contains one α-synuclein fold in either a singlet or doublet association [25, 32, 33].

The previously determined MSA filament types each consist of two asymmetric protofilaments (PF-A and PF-B) that associate along an extended interface nearly spanning the width of the filament core (Supplementary Fig. S1) [25]. Also common to each type is a central channel flanked by basic residues (Lys43, Lys45, and His50 of each protofilament) with unassigned density present within the channel. However, major conformational differences lie within the protofilaments that compose each mature fibril core. Type I MSA filaments contain two PFs, one PF-IA and one PF-IB. Residues Gly14 to Phe94 are resolved in PF-IA, and Lys21 to Gln99 are resolved in PF-IB. While both PF-IA and PF-IB contain a hairpin motif in the N-terminal region and a three-layered L-shaped motif in the C-terminal region, PF-IA also contains a single-layered L-shaped motif (residues 32–45) joining the N- and C-terminal motifs. Thus, the residues composing the terminal motifs differ in each protofilament, and the fold of the three-layered L-shaped motif also differs in the packing of the inner layer relative to the central layer.

The Type II filament subtypes (II_1_ and II_2_) each have two protofilaments [25] and contain identical PF-IIA, which span residues 14–94. Comparing PF-IA and PF-IIA, the N-terminal regions spanning residues 14–42 are conformationally identical. However, these protofilaments differ in the C-terminal region, including the conformation of the three-layered L-shaped motif. PF-IIA contains a small channel within the filament core, surrounded by Val52, Thr54, Ala56, Thr59, Glu61, Thr72, Gly73, and Val74, that is not present in PF-IIB because these residues form closer contacts with each other. PF-IIB_1_ and PF-IIB_2_ include residues 36–99 and contain a three-layered L-shaped motif (residues 47–99). In contrast to PF-IB, these protofilaments lack the N-terminal hairpin and contain a shorter one-layer L-shaped motif (residues 36–46) instead. PF-IIB_1_ and PF-IIB_2_ are distinguished from each other by a change in a surface loop at the C-terminal end of the ordered core (residues 81–90). The surface loop conformation of PF-IIB_1_ is similar to the loop in PF-IB, while PF-IIB_2_ has a distinct loop conformation that is shifted approximately 4 Å towards the C-terminus of the protofilament core. The overall helical symmetry of each MSA filament type is similar, although Type I filaments have a slightly greater twist (−1.42°) compared to Type II_1_/II_2_ filaments (−1.34°).

MSA filament types are heterogeneously distributed between different patients [25]. For example, in the original report from Schweighauser et al., two patients had almost exclusively Type I filaments, one patient had almost exclusively Type II filaments, and the remaining two patients had both Type I and II filaments in varying ratios. This heterogeneity suggests that there may be a greater variety of folds than previously reported. In this study, we use cryo-EM to obtain high-resolution (3.2–3.3 Å) structures of filaments purified from tissue of another patient with MSA. While the majority (66%) of the data consists of Type II_2_ filaments, our results reveal a novel protofilament fold in a subset (5%) of filaments that are related to Type I filaments.

## Methods

### MSA filament extraction

Deidentified human tissue was obtained from the Massachusetts Alzheimer’s Disease Research Center (ADRC). All donors provided consent to donate brains for research purposes in accordance with the standards of the ADRC. This study was exempt from institutional review board approval in accordance with the institutional review board policy of the University of California San Francisco. Filament extraction was performed using a modified protocol from Schweighauser et al. [25]. In summary, 1–2 g of freshly frozen cerebellum tissue from a patient pathologically diagnosed with MSA was homogenized at 20 mL/g of tissue in 10 mM Tris-HCl (pH 7.4), 800 mM NaCl, 1 mM EGTA, and 10% sucrose. N-lauroylsarcosinate (final concentration of 1% w/v) was added to the homogenate and incubated at 37°C for 30 min. Afterwards, homogenates were centrifuged at 10,000 × *g* for 10 min. The supernatant was kept and centrifuged at 100,000 × *g* for 20 min. The pellet was resuspended at 500 μL/g of frozen tissue in 10 mM Tris-HCl (pH 7.4), 800 mM NaCl, 1 mM EGTA, and 10% sucrose. The suspension was then centrifuged at 3,000 × *g* for 5 min. The supernatant was kept and diluted three-fold with 50 mM Tris-HCl (pH 7.4), 150 mM NaCl, 10% sucrose, and 2% sarkosyl. This suspension was then centrifuged at 150,000 × *g* for 1 h. The filament-enriched pellet was resuspended in 30 mM Tris-HCl (pH 7.4) using 100 μL/g of frozen tissue.

### Cryo-EM sample preparation and data collection

Purified filaments (3 μL) were added to a 200 mesh 1.2/1.3R Au Quantifoil grid coated with a 2-nm thick carbon layer, which was not glow discharged. After 30 seconds, grids were blotted for 7.5 s at room temperature and 100% humidity using a FEI Vitrobot Mark IV, followed by plunge freezing in liquid ethane. A total of 42,224 super-resolution movies were collected at a nominal magnification of 105,000× (physical pixel size: 0.417 Å/pixel) on a Titan Krios (Thermo Fisher Scientific) operated at 300 kV and equipped with a K3 direct electron detector and BioQuantum energy filter (Gatan, Inc.) set to a slit width of 20 eV. A defocus range of −0.8 to −1.8 μm was used with a total exposure time of 2.024 s fractionated into 0.025-second subframes. The total dose for each movie was 46 electrons/Å^2^. Movies were motion-corrected using MotionCor2 [34] in Scipion [6] and were Fourier cropped by a factor of 2 to a final pixel size of 0.834 Å/pixel. Motion-corrected and dose-weighted micrographs were manually curated in Scipion to remove micrographs lacking filaments, those at low resolution, or those with significant ice contamination, resulting in 4,392 remaining micrographs.

### Cryo-EM image processing

A graphical overview of the data processing workflow is provided in Supplementary Figure S2. All image processing was done in RELION 4 [14, 16]. Dose-weighted summed micrographs were imported into RELION 4. The contrast transfer function was estimated using CTFFIND-4.1 [23]. Filaments were manually picked, and segments were extracted with a box size of 900 pixels downscaled to 300 pixels, resulting in 257,982 segments. Reference-free 2D classification was used to remove contaminants and segments contributing to straight filaments, resulting in 255,032 remaining segments. These were re-extracted with a box size of 288 pixels without downscaling, followed by another round of reference-free 2D classification that did not filter out more contaminant segments. One round of 3D classification with image alignment was performed on the segments using a reference map consisting of the existing MSA Type I filaments (PDB code: 6XYO), a regularization parameter (T) of 20, and fixing helical parameters to −1.42° twist and 4.76 Å rise. We were unable to resolve Type I_2_ filaments by allowing the helical parameters to vary. One class (12,802 segments) corresponded to Type I_2_ filaments. One round of 3D auto-refinement was run using this map low-pass filtered to 10 Å, allowing rise and twist parameters to vary. The map was sharpened using the standard post-processing procedures in RELION. Full statistics are shown in Supplementary Table S1.

Two out of the 12 classes (57,809 segments) corresponded to Type II_2_ filaments with no breaks in the polypeptide density. Segments corresponding to these classes were pooled and subjected to one round of 3D classification without image alignment using a reference map consisting of one of these classes (A3; Supplementary Fig. S3), a regularization parameter (T) of 20, and fixing helical parameters to −1.34° twist and 4.76 Å rise. The highest resolution class (31,115 segments) was subjected to 3D auto-refinement and post-processing to give a refined Type II_2_ filament map.

Of the 12 classes, 4 classes (110,255 segments) displayed clear separation between protofilaments in the z-direction but modest resolution in the x- and y-directions, suggesting suboptimal alignment. Segments corresponding to these classes were pooled and subjected to one round of 3D classification with image alignment using a reference map consisting of the existing MSA Type II_2_ filaments (PDB code: 6XYQ), a regularization parameter (T) of 20, and fixing helical parameters to −1.34° twist and 4.76 Å rise. Most of these segments (110,071) converged into one class corresponding to Type II_2_ filaments, which was not further refined. The remaining classes (74,166 segments) from the initial round of 3D classification were low resolution and excluded from further processing.

To eliminate bias in 3D classification and further verify the distribution of filaments, we generated a 3D initial model ab initio from 2D class averages using RELION’s relion_helix_inimodel2d feature. This initial model was used as a reference map for 3D classification using the otherwise same inputs as the first round of 3D classification described above, resulting in classes corresponding to Type II_2_ filaments (206,869 segments, 81% of the data) and Type I_2_ filaments (15,369 segments, 6% of the data).

### Model building, refinement, and analysis

A single strand of the previously solved MSA Type II_2_ (PDB code: 6XYQ) or type I (PDB code: 6XYO) filament structures was placed in the density using Chimera [20]. Subsequent model building was performed using COOT [8] and ISOLDE [5] followed by refinement in Phenix [1]. The model was then translated to produce a stack occupying all regions with continuous density on the map. Refinement statistics are shown in Supplementary Table S1. Data were deposited into the Protein Data Bank (PDB) under accession codes 9CD9 (Type II_2_) and 9CDA (Type I_2_) and into the Electron Microscopy Data Bank (EMDB) under accession codes EMD-45464 (Type II_2_) and EMD-45465 (Type I_2_).

## Results

Partially purified, sarkosyl-insoluble filaments were isolated from the postmortem cerebellum of a 67-year-old male neuropathologically diagnosed with MSA. Negative-stain transmission electron microscopy on this sample revealed filaments that were morphologically similar to those in previous reports (Supplementary Fig. S4) [17, 25]. After cryo-EM imaging using standard collection methods (Fig. 1A; see Methods), reference-free 2D classification resulted in 2D class averages that were similar to those previously reported [25], exhibiting a crossover distance of approximately 600 Å (Supplementary Fig. S5). However, it was unclear whether these 2D classes corresponded to a known filament type. After 3D classification and helical reconstruction, we were able to obtain high-resolution structures of two MSA filament folds (Fig. 1B). The primary fold type (Fig. 1C), refined to 3.2 Å and representing 66% of the data (Supplementary Table S1), was nearly identical to Type II_2_ filaments (PDB: 6XYQ), with an all-atom root-mean-square deviation (RMSD) of 0.5 Å (Supplementary Fig. S6). We observed previously reported regions of unknown density, such as non-proteinaceous density within the central channel flanked by basic residues and a peptide-like density contacting residues Lys80–Val82 of PF-IIA [25]. We were able to resolve an additional residue (Leu100) at the C-terminal region of the PF-IIB_2_ ordered core.

**Fig. 1.**
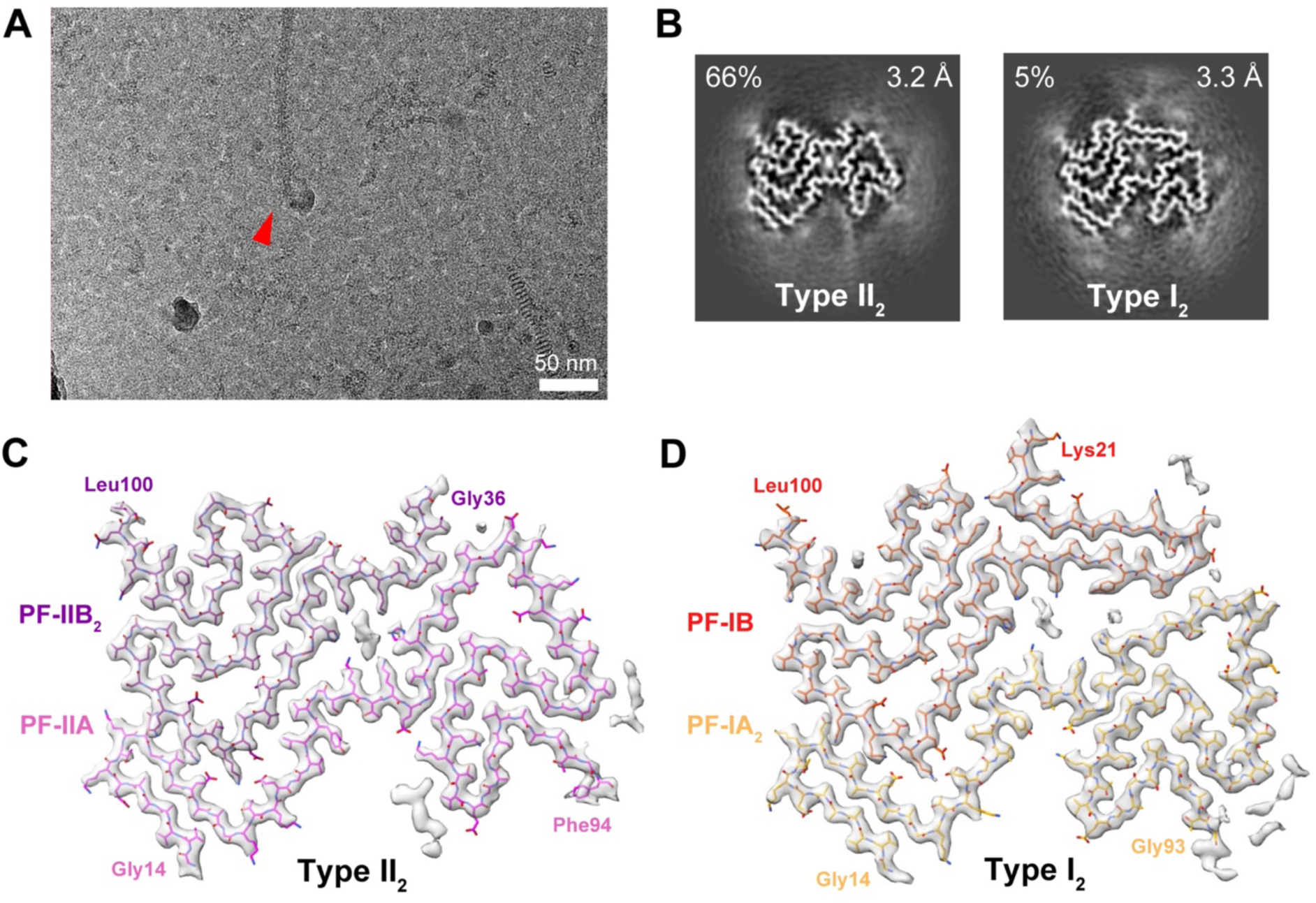
Cryo-EM analysis on ex vivo MSA filaments reveals a novel filament fold: Type I_2_ filaments. (**A**) Representative micrograph. The red arrow indicates an MSA filament. (**B**) Cross-section of two major conformations of MSA filaments, Type II_2_ and Type I_2_, identified after 3D classification of filament segments. The resolutions of the final reconstructions and abundance in the data are indicated. The remaining 29% of data consisted of low-resolution classes. (**C**) Density map and model of Type II_2_ filaments. The two constituent protofilaments PF-IIA and PF-IIB_2_, along with residues at the termini of the ordered core, are labeled and color coded in pink and purple, respectively. Regions of unmodeled density are also shown. (**D**) Density map and model of Type I_2_ filaments. The two constituent protofilaments PF-IA_2_ and PF-IB, along with residues at the termini of the ordered core, are labeled and color coded in red and yellow, respectively. Regions of unmodeled density are also shown.

In addition to the successful reproduction of the Type II_2_ MSA filament, we also resolved a filament exhibiting a conformation of α-synuclein that has not been reported previously. We refer to this low-populated filament fold as a Type I_2_ filament, which represents 5% of the data (Fig. 1B and Supplementary Fig. S7). The structure of Type I_2_ filaments, refined to 3.3-Å resolution, resembles Type I (hereafter called Type I_1_) and comprises two protofilaments, PF-IA_2_ and PF-IB (Fig. 1D). The ordered core of these protofilaments includes residues Gly14–Gly93 for PF-IA_2_ and Lys21–Leu100 for PF-1B. The PF-IB protofilament is common between Type I_1_ and Type I_2_ filaments, with an RMSD of 0.8 Å to the existing Type I_1_ filament model (PDB: 6XYO) [25].

The differences between Type I_1_ and Type I_2_ filaments exist within the second protofilament: PF-IA for Type I_1_ and PF-IA_2_ for Type I_2_ (Fig. 2A). While the N-terminal regions comprising residues 14–56 are nearly identical between the two protofilaments, the C-terminal regions of the protofilaments (residues 57–93) adopt distinct conformations. Specifically, this region in PF-IA_2_ is shifted outwards and away from the interprotofilament interface. This conformational change shifts the register of the hydrophobic intraprotofilament interface containing residues Val74 and Ala76 by two residues and creates a channel within the filament core, resulting in a net loss of hydrophobic interactions (Fig. 2B). Additionally, potential interstrand polar contacts between the Glu61 sidechain and Gly73 backbone amide are lost. Interestingly, the C-terminal region of PF-IA_2_ is virtually identical to the analogous region in PF-IIA found within Type II MSA filaments, although PF-IIA contains the C-terminal Phe94 residue that was not well resolved in PF-IA_2_ (Fig. 2C). Comparing PF-IA and PF-IA_2_, the average Cα displacement of residues 57–93 is 4.8 Å but is 0.3 Å between PF-IIA and PF-IA_2_.

**Fig. 2.**
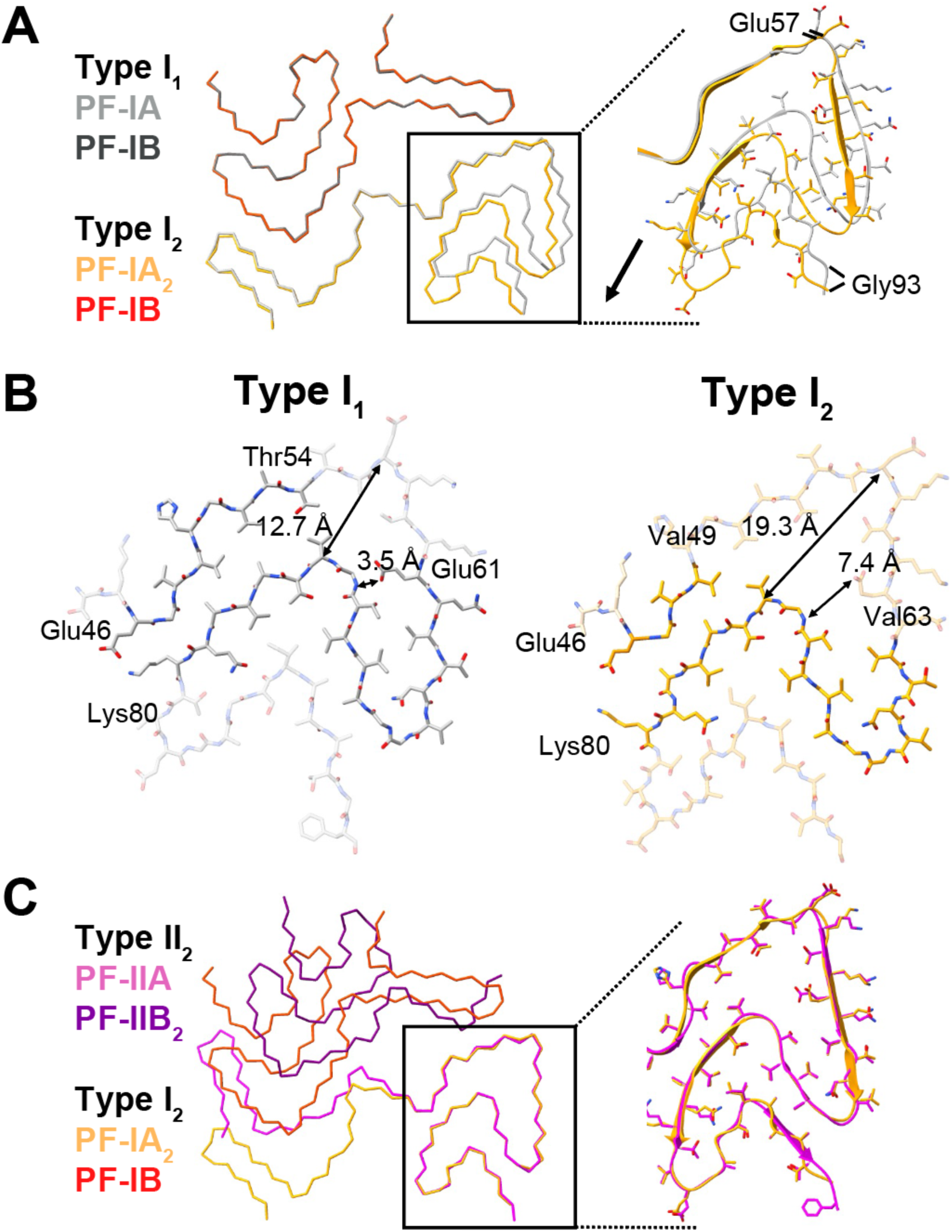
Comparison of MSA Type I_2_ filaments to previously solved MSA filament structures. (**A**) Overlay of Type I_1_ (PDB: 6XYO) and Type I_2_ filaments aligned using PF-IB. The black arrow indicates the direction of the conformational change in the C-terminal region. The average Cα displacement of residues 57–93 is 4.8 Å. (**B**) Comparison of intraprotofilament contacts in the C-terminal regions of PF-IA (Type I_1_) and PF-IA_2_ (Type I_2_). (**C**) Overlay of Type I_1_ and Type II_2_ (PDB: 6XYQ) filaments aligned using residues 47–93 of PF-IIA.

The morphological and helical parameter similarities between Type II_2_ and Type I_2_ MSA filaments prompted us to inquire whether both folds could coexist within the same filament or if the folds are instead segregated to individual filaments. Previous cryo-EM analysis of Lewy body α-synuclein filaments suggested that multiple morphologies (twisted and untwisted) could coexist within the same filaments based on 2D classification, although a structure of the untwisted morphology was not determined [33]. To address this question, we mapped segments classifying Type I_2_ or Type II_2_ conformations to the original micrographs (Fig. 3A–C). Due to the 13-fold greater abundance of Type II_2_ compared to Type I_2_ segments in the dataset, we only analyzed micrographs containing at least one Type I_2_ segment, corresponding to 437 micrographs out of 4,392 (10%). We found that 71% of the 556 filaments in these micrographs contained segments that classified exclusively or predominantly (90% or higher) to either Type I_2_ or Type II_2_ (Fig. 3D). The remaining minority of filaments contained a more mixed distribution of folds, although Type I_2_ tended to predominate (e.g., Fig. 3A). These results suggest that each MSA type preferentially segregates into separate filaments, though it is possible that a small number of filaments contain mixed types. A precise quantification of this dataset is complicated by the possibility of suboptimal classification, the extent of which is unknown.

**Fig. 3.**
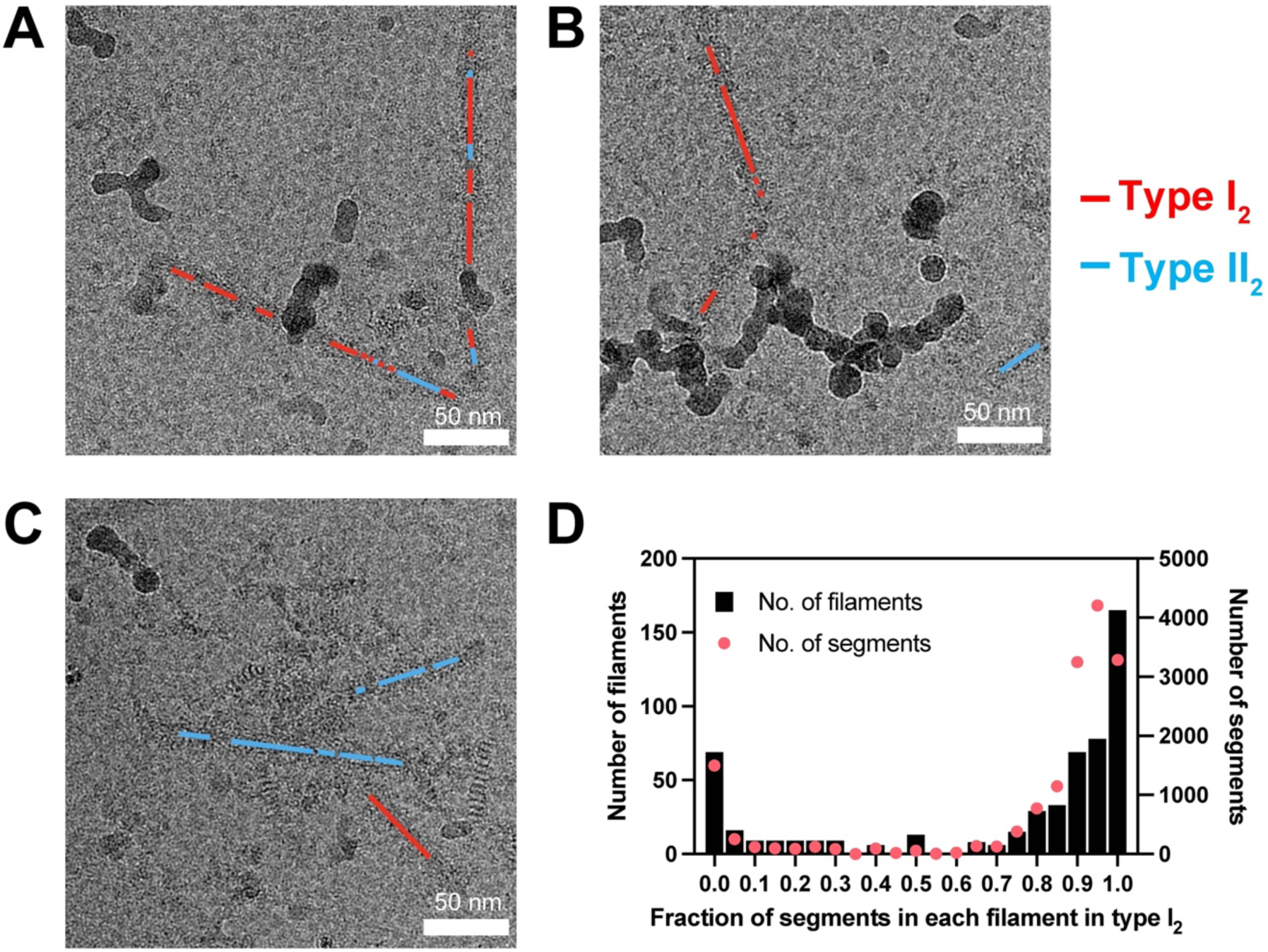
Analysis of MSA Type I_2_- and II_2_-classified segments within individual filaments. (**A–C**) After one round of 3D classification, segments corresponding to either type are color coded and mapped to the original micrographs. (**D**) Histogram of the fraction of segments classified as Type I_2_ per filament in micrographs containing at least one I_2_ segment.

## Discussion

Cryo-EM studies have been instrumental in determining the structures of disease-associated amyloid fibrils from synucleinopathies and other neurodegenerative diseases [9, 10, 13, 25, 27, 33]. Proteins associated with these diseases adopt numerous polymorphic folds that are associated with specific neurodegenerative diseases [24]. In some diseases, such as MSA, α-synuclein can adopt multiple distinct folds that are heterogeneously distributed between patients [25]. While Type I_1_ and II_2_ filaments appear to predominate in MSA, the lower-population type II_1_ also exists within some patients. Here, we have expanded the spectrum of known MSA filament types to include a fourth type, Type I_2_ filaments.

Type I_2_ filaments share one protofilament fold (PF-IB) with Type I_1_ filaments but differ in the PF-IA_2_ protofilament, which instead bears similarities with the C-terminal region of the PF-IIA protofilament found in Type II filaments. The chimeric nature of PF-IA_2_ suggests that common intermediates may exist in the misfolding and aggregation pathways of α-synuclein in MSA but then diverge, leading to multiple folds. The exact nature of these pathways and the relative contribution of each filament type to disease remains unknown. While MSA has two phenotypes—MSA-C (cerebellar) and MSA-P (parkinsonian) [11]—whether the identity and distribution of filament types correlates to phenotype remains undetermined.

We found that MSA Type I_2_ and II_2_ filaments tended to segregate to individual filaments, but it is possible that a small number of filaments contain a mixed population of both types. These results contrast with previous analyses of morphologically similar filament folds. Lewy body α-synuclein filaments were found to contain a roughly even distribution of both twisted and untwisted segments, although that analysis is limited by the lack of a structure of the untwisted morphology [33]. Another study investigating ex vivo antibody light chain filaments from a patient with systemic AL amyloidosis found two folds that unambiguously coexist in most filaments and differ only slightly in a surface-exposed region of the filament core [22]. The overall high homology allows for the formation of mixed filaments that maintain the favorable interstrand interactions needed for filament growth and stability. In the case of MSA Types I_2_ and II_2_, the folds are mostly dissimilar from each other except for residues 57–93 of PF-A, likely preventing the formation of a stable interface between both types. We attempted to model one Type I_2_ filament rung directly adjacent to a type II_2_ filament rung, which were aligned using the homologous residues 57–93 of PF-IIA (Supplementary Fig. S3). However, the model revealed severe steric clashes in the nonhomologous regions of the filaments. We do note the presence of a small number of filaments (11/556 filaments) containing numerous segments (≥10) classified into each type, suggesting that some mixed filaments may exist. It is possible that unresolved intermediate folds stabilize the interface between different fibrillar forms.

Overall, these results improve our understanding of the α-synuclein structures underlying MSA and provide a framework for developing therapeutics or diagnostics that bind to MSA filaments with high affinity and selectivity. The existence of multiple MSA α-synuclein folds suggests that these putative molecules should target a conserved region of the filament. However, conformation-specific molecules would also be useful for rapidly quantifying each filament type in a tissue sample without needing to resort to cryo-EM [4, 26, 32]. These molecules should help us continue investigating the extent to which multiple folds exist in single filaments.

## List of Abbreviations

ADRC: Alzheimer’s Disease Research Center
cryo-EM: Cryo–electron microscopy
EMDB: Electron Microscopy Data Bank
FSC: Fourier shell correlation
GCI: Glial cytoplasmic inclusions
MSA: Multiple system atrophy
PDB: Protein Data Bank
PF: Protofilament
RMSD: Root-mean-square deviation

## Declarations

### Ethics approval and consent to participate

Not applicable

### Consent for publication

Not applicable

### Availability of data and materials

Refined atomic models have been deposited in the Protein Data Bank (PDB) under accession numbers 9CD9 (Type II_2_) and 9CDA (Type I_2_). Corresponding cryo-EM maps have been deposited in the Electron Microscopy Data Bank (EMDB) with accession numbers EMD-45464 (Type II_2_) and EMD-45465 (Type I_2_). Please address requests for materials to the corresponding author.

### Author Contributions

N.Y., D.R.S., N.A.P, and G.E.M designed the research; N.Y., F.C., E.T., A.M., D.A.M., and G.E.M performed experiments; N.Y., S.B.P., D.R.S., N.A.P., and G.E.M. analyzed the data; N.Y., D.R.S., N.A.P., and G.E.M. wrote the manuscript. All authors read and approved the final manuscript.

## Acknowledgements

We thank the patients’ families for donating brain tissue.

---

**Supplementary Table S1.** Cryo-EM data collection and model refinement statistics.

| **Data collection and processing** | **MSA Type II<sub>2</sub>** | **MSA Type I<sub>2</sub>** |
| --- | --- | --- |
| Microscope and camera | Titan Krios, K3 |  |
| Magnification | 105,000 |  |
| Voltage (kV) | 300 |  |
| Electron exposure (e<sup>-</sup>/Å<sup>2</sup>) | 46 |  |
| Dose rate (e<sup>-</sup>/physical pixel/sec) | 16 |  |
| Exposure per frame (sec) | 0.024 |  |
| Defocus range (μm) | -0.8 to -1.8 |  |
| Physical pixel size (Å) | 0.834 |  |
| Movies collected | 42,224 |  |
| Box size (pixels) | 288 |  |
| Interbox distance (Å) | 28 |  |
| Initial segments extracted | 257,982 |  |
| Final segments | 31,115 | 12,802 |
| Resolution (Å) | 3.2 | 3.3 |
| B-factor (Å<sup>2</sup>) | -73.0 | -74.8 |
| Helical rise (Å) | 4.76 | 4.76 |
| Helical twist (°) | -1.35 | -1.42 |
| **Refinement** | **MSA Type II<sub>2</sub>** | **MSA Type I<sub>2</sub>** |
| **Model composition** |  |  |
| Non-hydrogen atoms | 10,950 | 10,989 |
| Protein residues | 1,600 | 1,606 |
| Number of chains | 20 | 22 |
| **R.M.S. deviations** |  |  |
| Bond lengths (Å) | 0.006 | 0.006 |
| Bond angles (°) | 0.586 | 0.530 |
| **Validation** |  |  |
| MolProbity score | 2.00 | 1.50 |
| Clashscore | 7.78 | 9.34 |
| Rotamer outliers (%) | 1.94 | 1.01 |
| Cβ outliers (%) | 0 | 0 |
| **Ramachandran plot** |  |  |
| Favored (%) | 94.87 | 98.59 |
| Allowed (%) | 5.13 | 1.41 |
| Outliers (%) | 0 | 0 |
| **PDB accession code** | **9CD9** | **9CDA** |
| **EMDB accession code** | **EMD-45464** | **EMD-45465** |

**Supplementary Fig S1.**
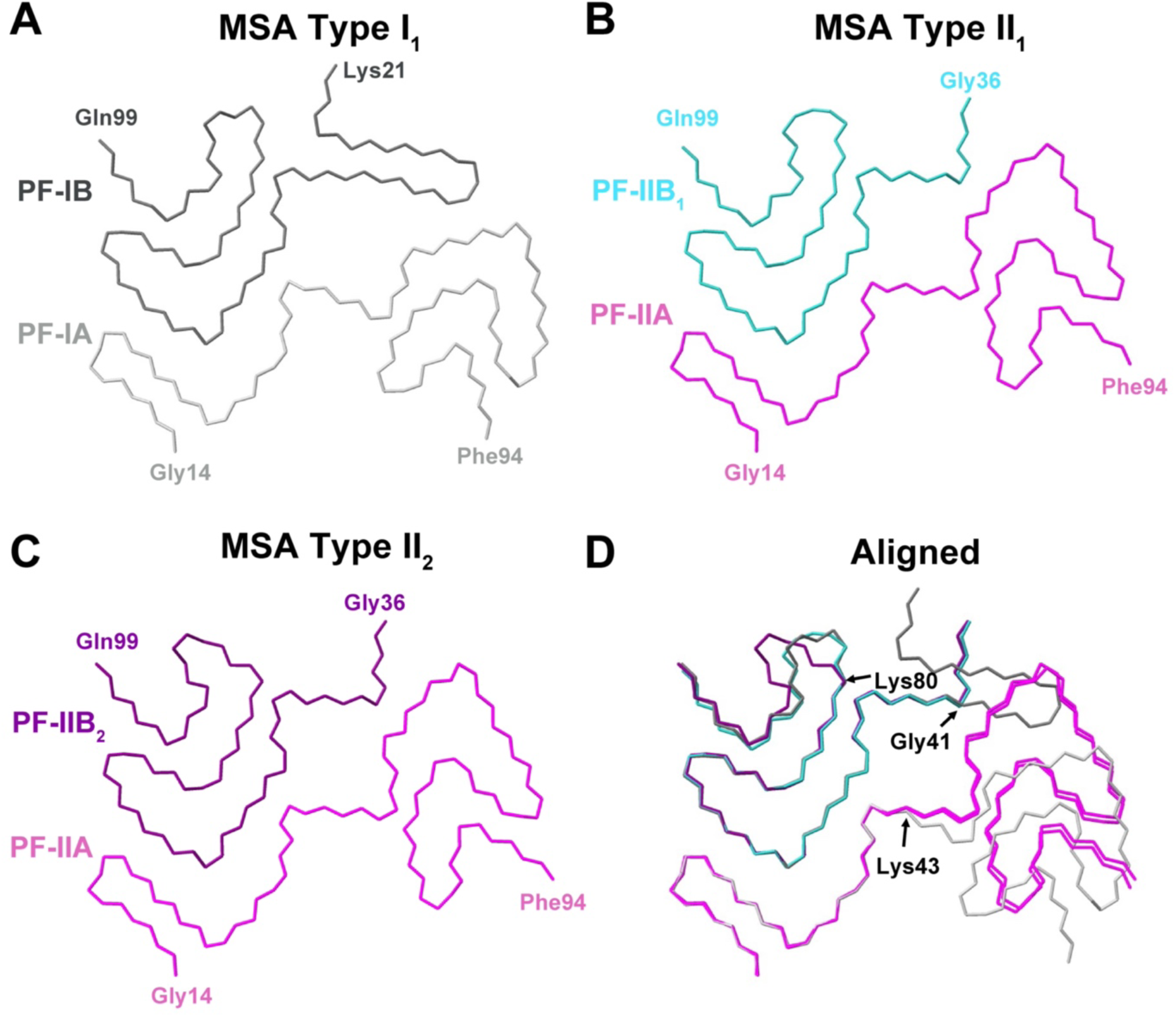
Structures of MSA filaments reported by Schweighauser et al. (**A**) Type I_1_ (PDB: 6XYO), (**B**) Type II_1_ (PDB: 6XYP), and (**C**) Type II_2_ (PDB: 6XYQ). (**D**) Alignment of the three filament conformations. In (**A**–**C**), residues at the termini of the ordered region of each protofilament are labeled. In (**D**), residues at the start or end of regions where the filament conformations diverge are labeled.

**Supplementary Fig S2.**
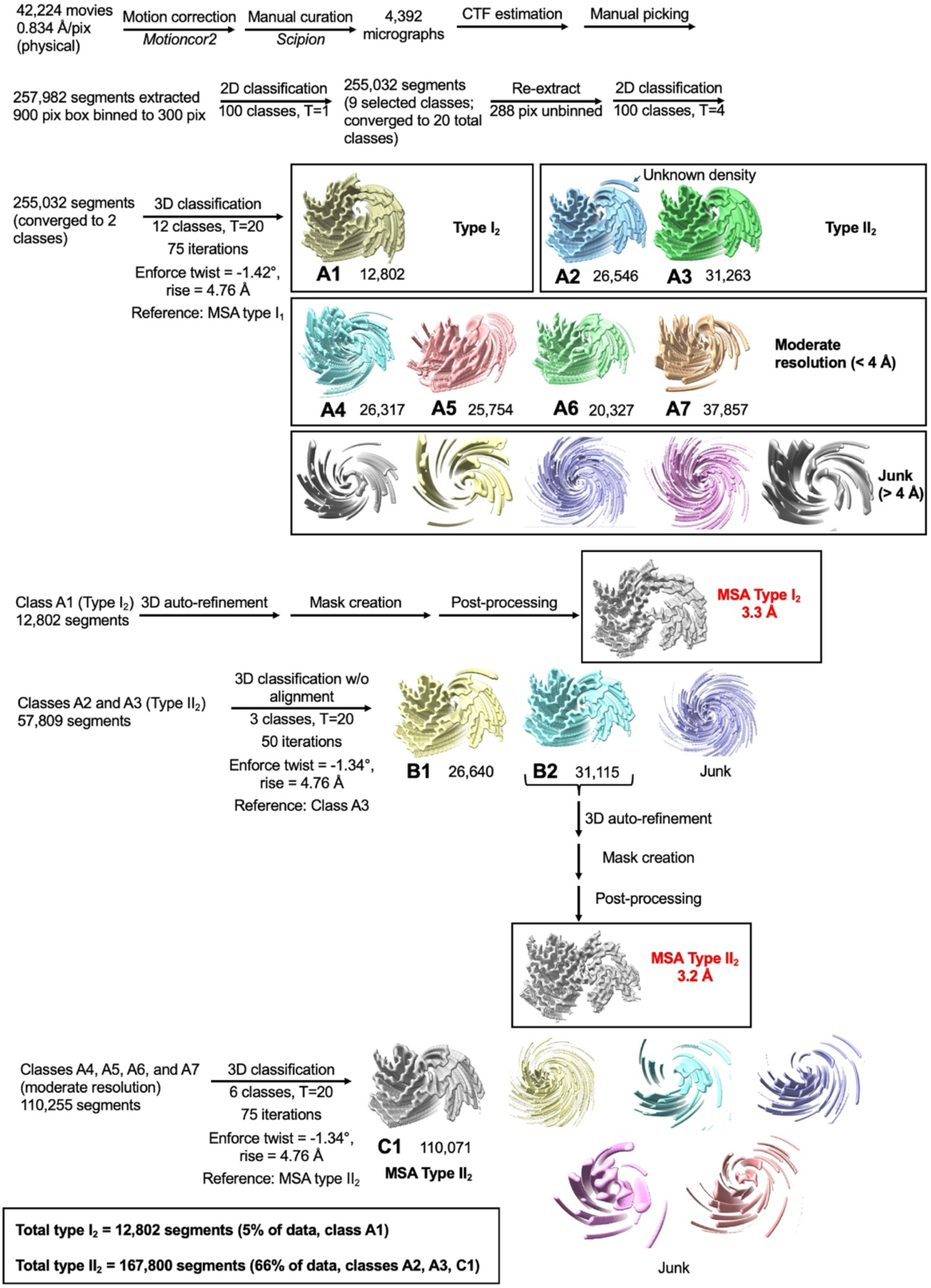
Flowchart of MSA filament data processing. Unless otherwise indicated, all data processing procedures were performed in RELION 4.

**Supplementary Fig S3.**
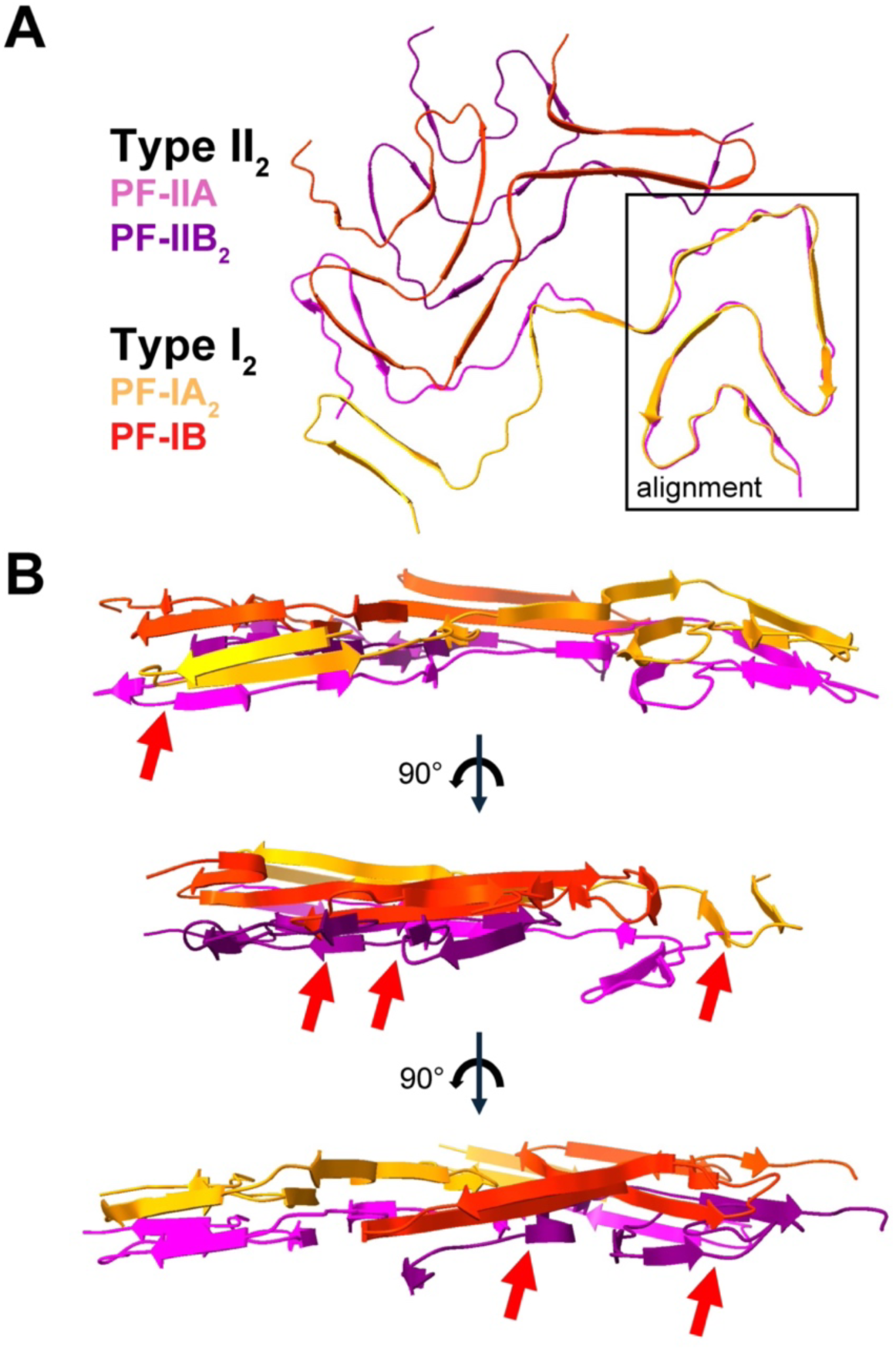
Model of MSA Type II_2_-I_2_ mixed cofilament wherein adjacent Type II_2_ and Type I_2_ rungs were aligned using residues 57–93 of PF-IIA (inset). (**A**) Cross-section of model. (**B**) Side-on views demonstrating severe steric clashes (red arrows) between adjacent protofilaments.

**Supplementary Fig S4.**
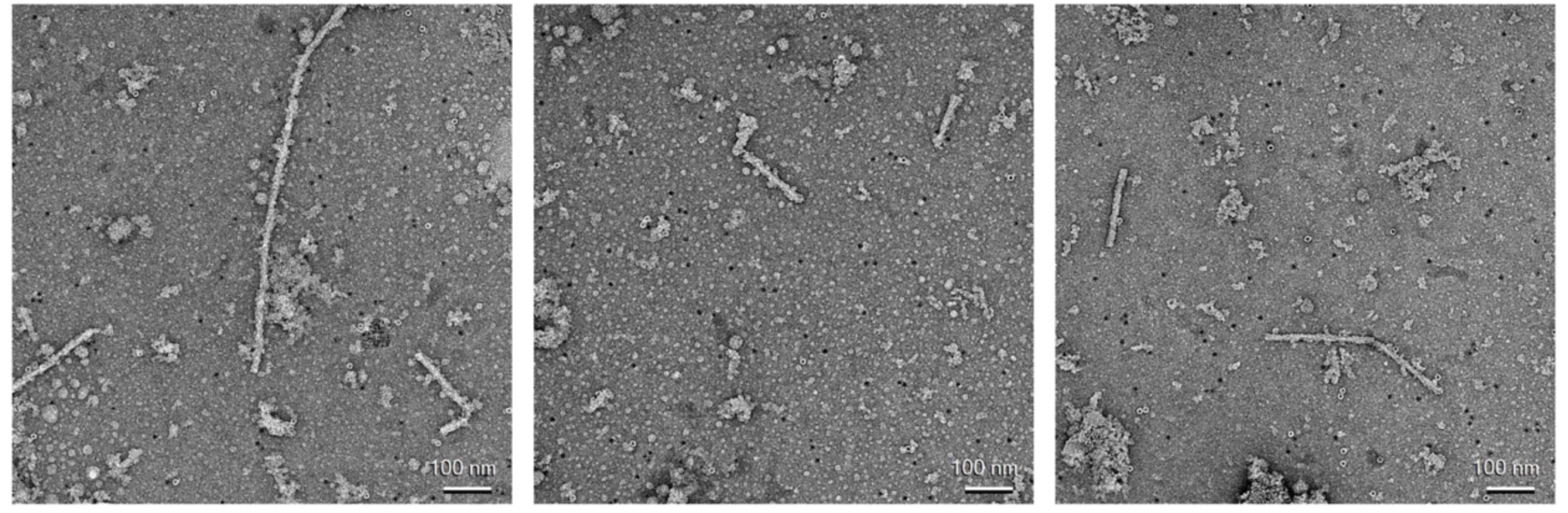
Representative negative-stain transmission electron micrographs of ex vivo MSA filaments.

**Supplementary Fig S5.**
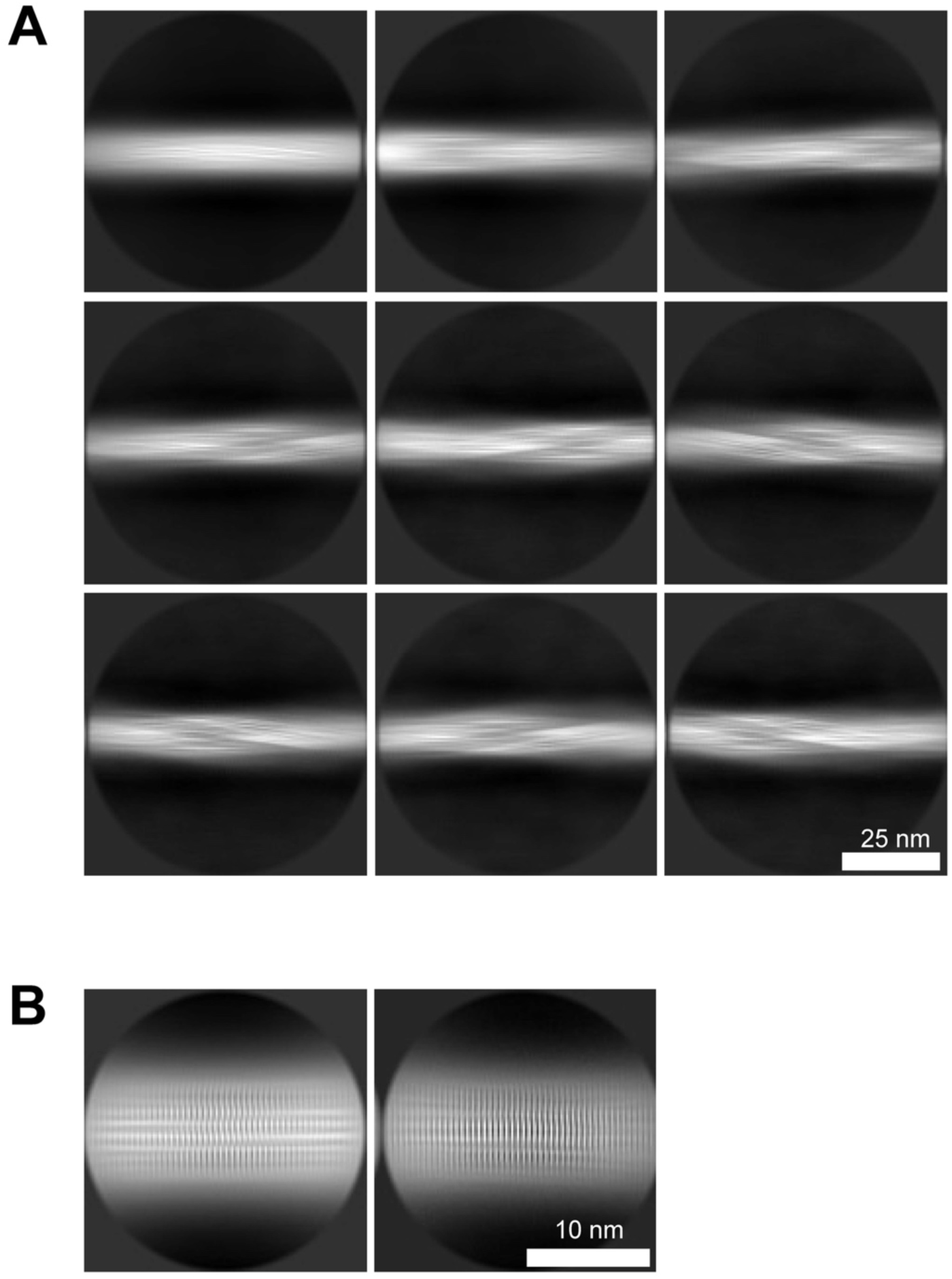
Two-dimensional class averages of MSA filaments extracted at two different box sizes. (**A**) At 900 pixels downscaled to 300 pixels. (**B**) At 288 pixels. Segments corresponding to these classes were used for 3D classification.

**Supplementary Fig S6.**
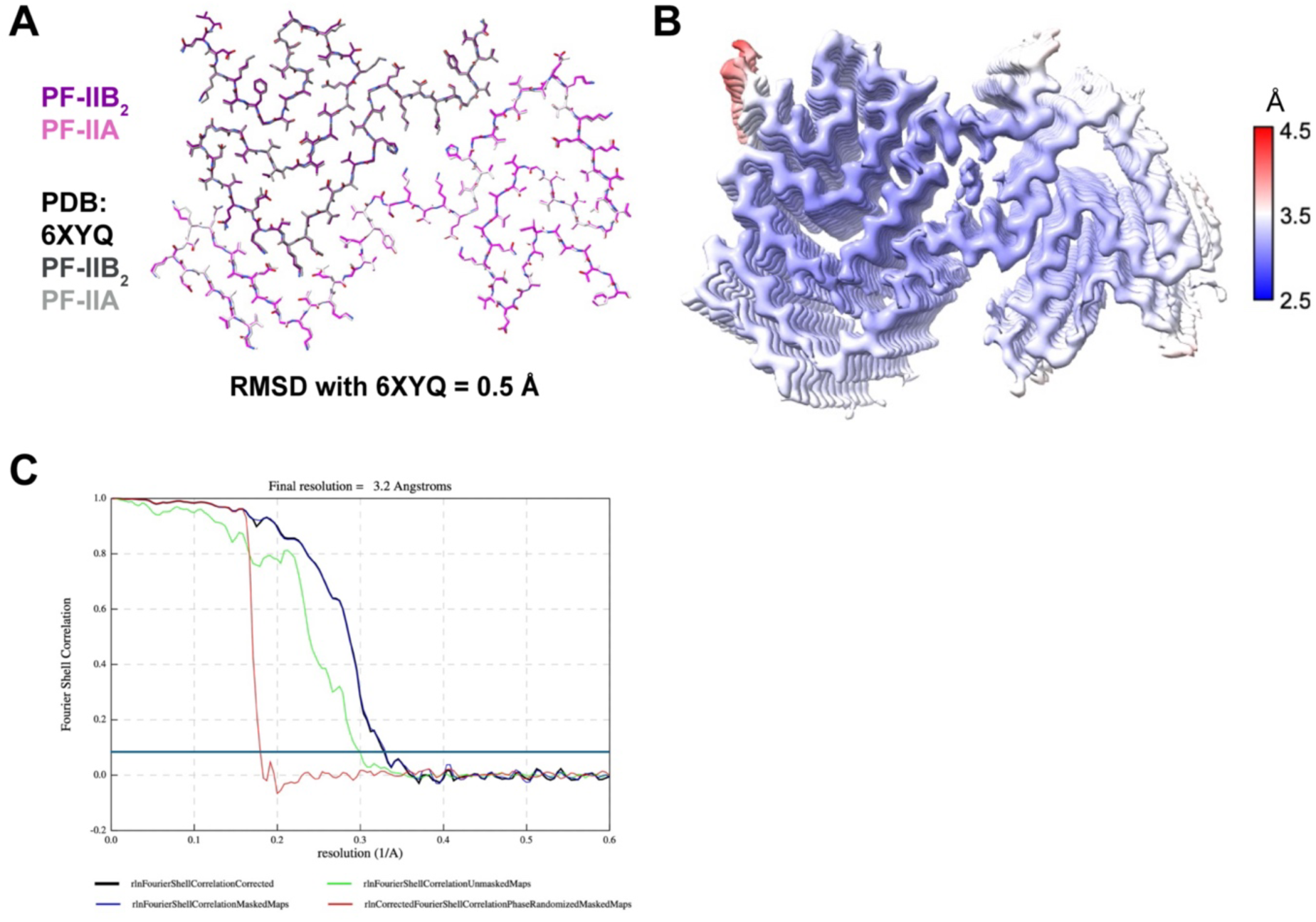
Characterization of MSA Type II_2_ filaments in the dataset. (**A**) Alignment of our Type II_2_ filament model with the previously published model (PDB: 6XYQ). (**B**) Local resolution of the Type II_2_ filament map. (**C**) Fourier shell correlation (FSC) curves of the Type II_2_ filament map. The dark blue line denotes the FSC = 0.143 cutoff used for resolution estimation.

**Supplementary Fig S7.**
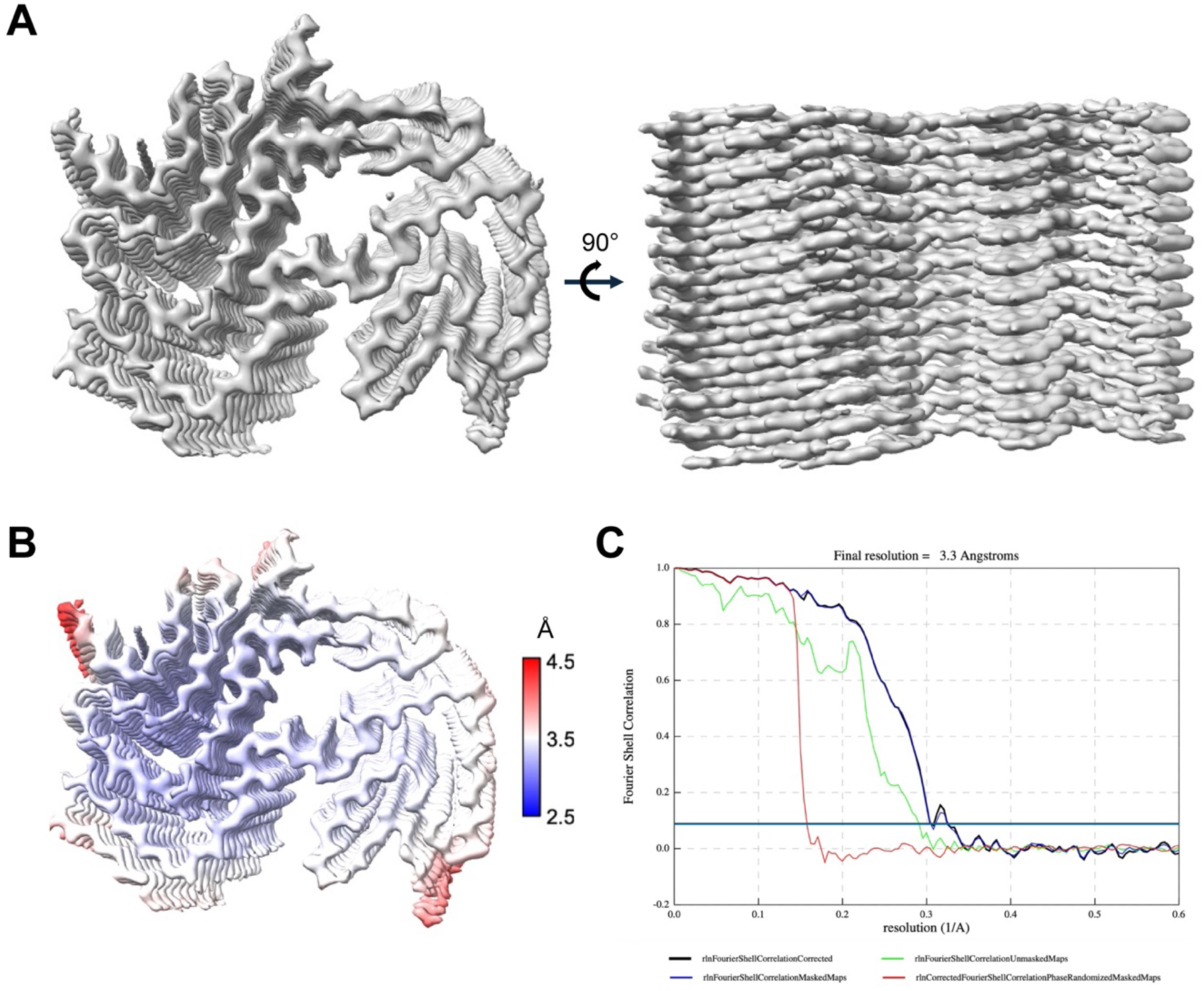
Characterization of MSA Type I_2_ filaments in the dataset. (**A**) Density map of Type I_2_ filaments showing separation between protofilaments in the long axis of the filament. (**B**) Local resolution of the Type I_2_ filament map. (**C**) FSC curves of the Type I_2_ filament map. The dark blue line denotes the FSC = 0.143 cutoff used for resolution estimation.

